# Investigating the basis of lineage decisions and developmental trajectories in the dorsal spinal cord through pseudotime analyses

**DOI:** 10.1101/2023.07.24.550380

**Authors:** Sandeep Gupta, Eric Heinrichs, Bennett G. Novitch, Samantha J. Butler

**Author notes:** Correspondence, CHS South Tower 67-200J, 650 Charles E Young Drive East, University of California, Los Angeles, Los Angeles CA 90095. Equal contributions (Sandeep Gupta, Eric Heinrichs).

## Abstract

Dorsal interneurons (dIs) in the spinal cord encode the perception of touch, pain, heat, itch, and proprioception. While previous studies using genetic strategies in animal models have revealed important insights into dI development, the molecular details by which dIs arise as distinct populations of neurons remain incomplete. We have developed a resource to investigate dI fate specification by combining a single-cell RNA-Seq atlas of mouse ESC-derived dIs with pseudotime analyses. To validate this *in silico* resource as a useful tool, we used it to first identify novel genes that are candidates for directing the transition states that lead to distinct dI lineage trajectories, and then validated them using *in situ* hybridization analyses in the developing mouse spinal cord *in vivo*. We have also identified a novel endpoint of the dI5 lineage trajectory and found that dIs become more transcriptionally homogenous during terminal differentiation. Together, this study introduces a valuable tool for further discovery about the timing of gene expression during dI differentiation and demonstrates its utility clarifying dI lineage relationships.

**Summary statement:** Pseudotime analyses of embryonic stem cell-derived dorsal spinal interneurons reveals both novel regulators and lineage relationships between different interneuron populations.

## Introduction

Somatosensation permits us to perceive touch, temperature, pain (nociception), and hold our bodies correctly in space (proprioception). These sensory modalities are critical for daily life, as well as emotional well-being. Sensory information is received in the periphery and transmitted to higher-order centers in the brain, or spinal motor circuits, by sensory relay circuits in the dorsal spinal cord (Lai et al., 2016). These circuits arise from six populations of dorsal interneurons (dI1-dI6) with distinct molecular signatures, connectivity, and sensory functions (Andrews et al., 2017; Gupta and Butler, 2021). dIs emerge during embryonic development in response to multiple patterning and differentiation signals. Previous studies have shown that the bone morphogenetic protein (BMPs) and Wnt families act from the roof plate at the dorsal midline to direct the dorsal-most dI identities (dI1-dI3) (Andrews et al., 2017; Gupta et al., 2022; Hazen et al., 2012; Le Dreau et al., 2012; Lee et al., 1998; Liem et al., 1997; Liem et al., 1995; Megason and McMahon, 2002; Muroyama et al., 2002; Yamauchi et al., 2008). The fate specification process for the intermediate dI identities (dI4-dI6) is less well defined, but a recent study showed that retinoic acid (RA) is sufficient to direct these fates *in vitro* (Gupta et al., 2022), suggesting a role for RA acting from the paraxial mesoderm *in vivo*. Once differentiated, dIs then migrate to the correct laminae in the adult dorsal horn to form distinct sensory circuits (Koch et al., 2018).

The genetic program that directs dI differentiation remains unresolved. Key transcription factors have been identified that promote dorsal progenitor (dP) and dI identities (Lai et al., 2016), which include Atoh1, Neurog1/2, Ascl1 and Ptf1a. These factors are necessary and sufficient to specify some of the dI populations (Bermingham et al., 2001; Glasgow et al., 2005; Gowan et al., 2001; Helms et al., 2005; Mizuguchi et al., 2006; Wildner et al., 2006). However, it has remained unclear how a limited number of growth factors regulate these transcription factors to specify six distinct dI populations. We are assessing these mechanisms by developing embryonic stem cell (ESC) models (Andrews et al., 2017; Gupta et al., 2018; Gupta et al., 2021); most recently, we have described an improved protocol which can generate the complete complement of dIs with the correct functional and molecular signatures (Gupta et al., 2022). Stem cell models offer many advantages for mechanistic discovery including an unparalleled ability to control growth conditions and probe cellular/molecular responses in large populations of synchronously developing cells without the confounding effects of embryonic redundancy and lethality (Gaspard and Vanderhaeghen, 2010; Veenvliet et al., 2021; Zhu and Huangfu, 2013). Our studies using these models have suggested that BMPs do not establish dI fate by acting as morphogens (Andrews et al., 2017), dI fates rather appear to be established in a series of nested choice points (Gupta et al., 2022). Spinal progenitors are initially dorsalized by RA, subdivided into multipotential dP subgroups by RA±BMP signaling, and then directed into specific dI fates by as yet unknown mechanisms.

Here, we leverage our previously acquired single-cell (sc) RNA-Seq atlas of mESC-derived dIs (Gupta et al., 2022), to develop a tool to identify novel genes that are associated with dI fate specification. This tool combines the scRNA-Seq atlas with pseudotime analyses to reconstruct dI-specific lineage trajectories. This mESC-derived dI atlas was then compared to an scRNA-Seq mouse embryonic spinal cord dataset (Delile et al., 2019), to assess the similarities between lineage trajectories. There was generally broad concurrence about the lineage relationships, although some details differed. To identify the transitional states that precede the key choice points, we identified candidate genes *in silico*-which were then validated *in vivo* by analyses of their expression patterns in the developing mouse spinal cord. Our studies also investigated the endpoint of the dI5 lineage trajectory, and the emergence of distinct dI5 subtypes. We further observed that the paths of the different dI s converge upon terminal differentiation both *in vivo* and *in vitro*, i.e., dIs assume neuronal identities which are more transcriptionally similar than during their preceding developmental trajectories. Taken together, this analysis provides further understanding of dI lineage relationships and develops a resource for identifying novel developmental regulators of sensory circuit formation.

## Material and Methods

### Seurat data processing and integration

Cellranger output (Gupta et al., 2022) was loaded into R (4.1.3) (R Core Team, 2022) and Seurat (Hao et al., 2021) (v4.0.4 - v4.1) and separate objects were made for the RA and RA+BMP protocols with min.cells = 3 and min.features = 200. Both datasets were then filtered for quality control based on violin plots of metadata with the goal of removing outliers on both ends (RA: nFeature > 2500, nCount_RNA > 5000 and < 50000, percent.mito < 10; BMP: nFeature > 200, nCount_RNA > 5000 and < 35000, percent.mito < 7). SCTransform (V1) was run on both datasets individually. Standard dimensional reduction was followed with 40pcs, three UMAP dimensions, and default cluster resolution.

To isolate cell types relevant to the desired differentiation and to remove unwanted byproducts and low-quality cells, the data were then subsetted to include only *Sox2*^+^ or *Tubb3*^+^ clusters that were also *Nanog*^-^ (pluripotent stem cells) and *Sox10*^-^ (neural crest cells), to remove cell types that were unrelated to the differentiation (Fig. S1A, B), and reprocessed with SCTransform pipeline. The two datasets were next integrated in Seurat (v4.2) using 3000 integration features and reciprocal principal component analysis (RPCA) based on principle components (PCs) calculated from commonly varying genes rather than a canonical correlation analysis to avoid overfitting. The combined data were dimensionally reduced and embedded into three UMAP dimensions using 40 PCs. Clustering was performed using a resolution of 2 to obtain clusters that roughly correlated with differentiation trajectory and timepoint in differentiation. PrepSCTFindMarkers was run to correct counts from different datasets to aid in further expression analysis. All plots were generated using ggplot2 (Wickham, 2016), Plotly (Fahd Qadir, 2019; Qadir et al., 2020; Sievert, 2020), and Graphpad Prism.

### Monocle pseudotemporal ordering

To find pseudotime trajectories in our combined dataset, we transferred our data to Monocle3 (Cao et al., 2019) using SeuratWrappers, and the cells were clustered using the UMAP reduction and Learn_Graph run to ascertain the principal graph of the data. The parameters were optimized to close the overall loop of the dataset and provide sufficient branching without yielding erroneous branches (use partition = F, learn_graph_control: Euclidian distance ratio = 2, geodesic distance ratio = 1/5, minimal branch length = 10, orthogonal project tip = F, n_center = 340, prune graph = T). Choose_cells was used to select all cells in the progenitor area up to the initial bottleneck and these were all set as the root when running order_cells. These data were then added back into Seurat as a metadata column to be accessed during further analysis.

### Pseudotime and marker gene analysis

Marker gene analysis was done with Seurat v4.3. FindAllMarkers (only.pos = T) was run on the combined dataset to identify marker genes for each cluster. The Seurat object was split into 5 sub-objects, one for each of the 5 trajectories, by subsetting on clusters. To analyze expression over pseudotime, Locally Estimated Scatterplot Smoothing with a span of 0.3 was used to create a curve that fit the data and predict gene expression at every 0.1 pseudotime value. These values were then plotted in ggplot2 using geom_tile.

Analysis of the different pseudotime timepoints was achieved by splitting the data into progenitor (the cells used as the root in Monocle), dP (cells less than 7.5 Pseudotime), or dI (cells greater than or equal to 7.5 Pseudotime) groupings. The cutoff between dP and dI was assigned based on when expression of most dP markers peaked, and where secondary splits in the trajectories occurred in the UMAP expression plots. FindAllMarkers was run and any positive marker gene with an adjusted p value < 0.05 was submitted to Metascape (Zhou et al., 2019) for gene ontology (GO) and other analyses. Similarly, *Sncg* positive cells were analyzed by taking cells in clusters 16 or 7 with a *Sncg* expression value greater than 1 (and a Pseudotime > 0.3 due to a clustering issue) and running FindMarkers against the remaining cells. Metascape-based plots were made in Prism.

### Analysis of the in vivo spinal cord dataset

#### Construction of the *in vivo* atlas

*In vivo* data was downloaded from https://www.ebi.ac.uk/biostudies/arrayexpress/studies/E-MTAB-7320. Ensembl IDs were converted to MGI symbols using BioMart, IDs without a corresponding symbol, or symbols aligning to multiple IDs, were left out. This latter category included the genes *Bfar*, *Pakap*, *Gm16364*, *Gm16701*, *2933427D14Rik*, and *Gm16499*. The data were processed in Seurat and cleaned (nFeature_RNA > 1000, nFeature_RNA < 7500, percent.mt < 6, nCount_RNA < 40,000), and then subsetted to include only progenitors and neurons as defined by (Delile et al., 2019). The data were then split by timepoint, SCTransformed, and reintegrated in Seurat using rPCA integration. The same pipeline was used as above, i.e., 40 PCs and three components for UMAP, and 30 PCs for FindNeighbors. The Z component of the UMAP was inverted to visually align with the *in vitro* dataset. PrepSCTFindMarkers was run to correct gene expression. Cell identities were assigned using the most highly expressed gene in each cell, with ties broken at random.

#### Integration of the *in vivo* and *in vitro* datasets

The two cleaned datasets were combined into the list of datasets before SCTransform and integration, and same pipeline used as above. To assess similarity between UMAPs, UMAP coordinates assigned by us, and cell identities assigned by (Delile et al., 2019) were used to train a k nearest neighbor classifier with a k of 10. This classifier was then used to predict identities in our dataset using the integrated UMAP coordinates.

### In situ hybridization and immunohistochemistry

Digioxigenin (DIG)-labeled RNA probes against the 3′ untranslated regions of genes of interest were generated using the Roche RNA Labeling Kit and hybridized onto 12-14μm transverse sections of embryonic spinal cords. *In situ* hybridization (ISH) signals were visualized using anti-DIG antibody conjugated with an alkaline phosphatase fragment (Roche) and nitro-blue tetrazolium and 5-bromo-4-chloro-3’-indolyphosphate substrates. Target sequences were amplified using cDNA derived from the mouse embryonic stem cell (mESC)-derived spinal cord cell types using the primers listed in Table 1. All primers were designed with the Primer 3 program (http://primer3plus.com/) and T7 promoter sequence was added on all the reverse primers for generating antisense mRNA probes using T7 RNA polymerase (Roche). For immunohistochemistry (IHC) spinal cord sections were directly treated with 1% antibody blocking solution (1% heat inactivated horse serum in 1xPBST) for 1 hour at room temperature followed by incubation with the primary antibodies overnight at 4°C. The slides were then processed for ISH using standard techniques (Vesque et al., 2000). Fluorescently labelled species-specific secondary antibodies (Jackson Immunoresearch Labs) were used to detect the signal. The following primary antibodies were used: Lhx2 (goat, 1:250, Santa Cruz Biotechnology, catalog number: sc-19344), Lmx1b (guinea pig, 1:100, gift from Thomas Mueller, Dresden), Pax2 (rabbit, 1:500, Invitrogen, catalog number: 71-6000), Sncg (rabbit, 1:500, LSBio, catalog number LS-B14232-50). Sections were then counterstained with DAPI and imaged on a Zeiss LSM800 confocal system.

All animals were housed within controlled access facilities and were under the care and supervision of animal care technicians supervised by the UCLA veterinarians of the Division of Laboratory Animal Medicine. Permission for animal experimentation was granted by the UCLA Institutional Animal Care and Use Committee.

## Results

### Creation of a single cell atlas with the full repertoire of dorsal progenitors, transition states, and dorsal interneurons

To develop a resource to identify novel genes that direct neural fate specification and differentiation in the dorsal spinal cord, we mined our previously established single cell (sc) transcriptomic dataset that represents a complete atlas of *in vitro*-derived dorsal interneurons (dI) (Gupta et al., 2022). In brief, mESCs were converted into posterior neuromesodermal progenitors (NMPs) though the addition of basic FGF and the GSK3β antagonist CHIR 99021 (CHIR) (Gouti et al., 2014). These NMPs were then differentiated into dIs through the addition of either retinoic acid (RA) or RA together with bone morphogenic protein (BMP) 4 (Fig. 1A). These two protocols respectively generate either dorsal progenitors (dP)4 - dP6 or dP1-dP3, which then give rise to mature dI4-dI6s (RA protocol) or dI1-dI3s (RA+BMP4 protocol). These heterogenous cell populations were collected at day 9 of differentiation and processed for scRNA-Seq (Fig. 1A). Downstream analyses were performed to first compile an *in vitro*-derived single cell atlas for the dIs (Gupta et al., 2022), and then perform pseudotemporal ordering to identify candidate genes that direct fate changes at transition points (Fig. 1B, see also methods).

**Figure 1:**
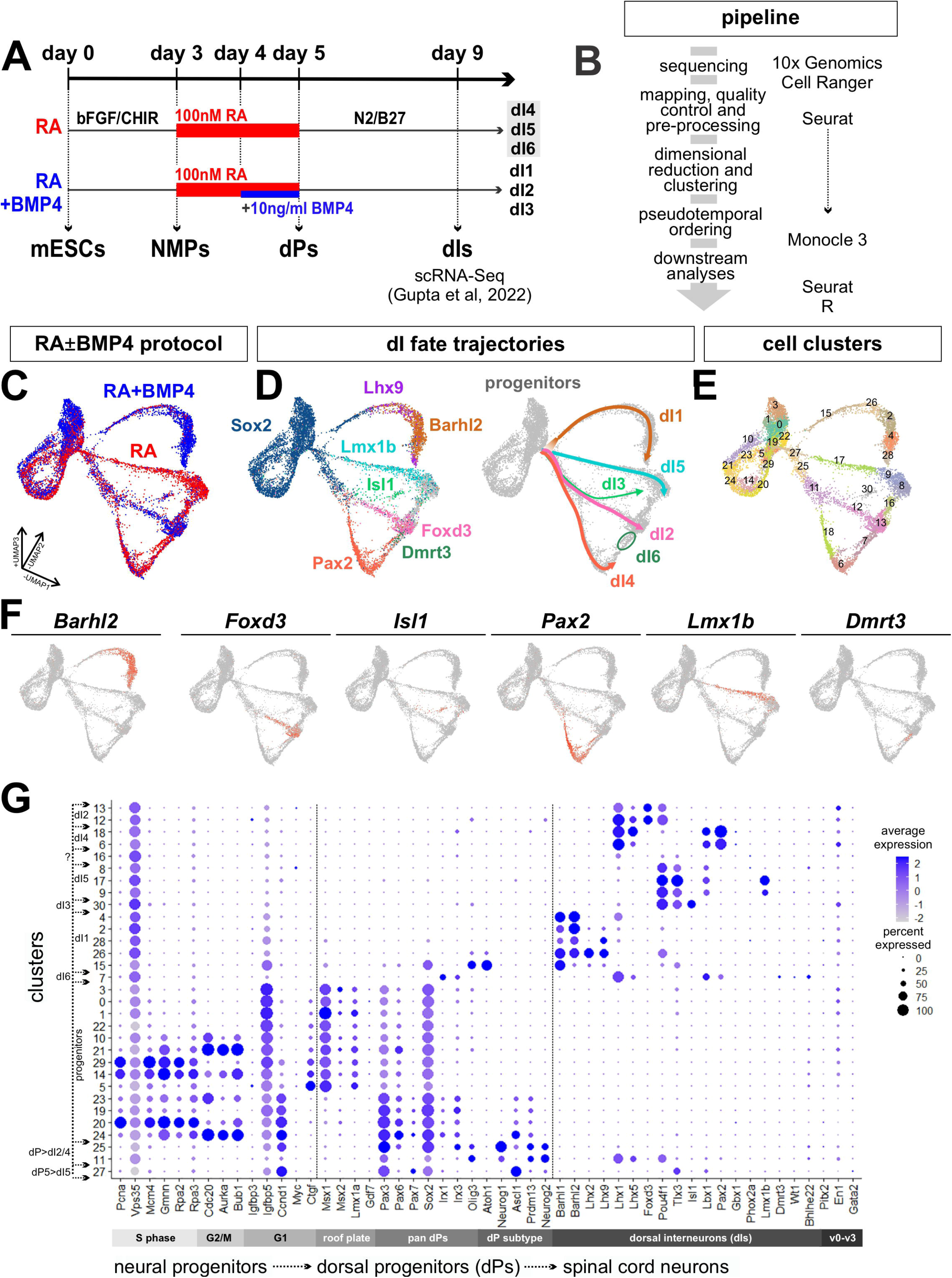
Single-cell analysis pipeline to identify dorsal interneuron lineages. (A) Schematic timeline for the derivation of dorsal interneurons (dIs) from mouse embryonic stem cells (mESCs). On day 9 of the differentiation, cells were dissociated and subjected to single-cell RNA sequencing (Gupta et al., 2022). (B) Overview of the pipeline for analysis of the single-cell transcriptomic data. (C-E) UMAP plots depict the combined cell types derived through the RA±BMP4 protocols (C), and the distinct dI lineages, as designated by marker analysis (D). Unsupervised clustering results in 31 distinct transcriptional clusters (E). (F) UMAP feature plots showing the expression of cardinal markers for all six classes of dIs. (G) Dot plot analysis showing the expression of various genes, which groups clusters in a continuous stream of progenitors to dorsal sensory interneurons (dIs).

Projection of both datasets using Seurat (Hao et al., 2021) into the same three-dimensional UMAP space reveals the overlap and divergence in the cell types arising from the RA (red) and RA+BMP4 (blue) protocols (Fig. 1C). The datasets generally overlap in the *Sox2*^+^ progenitor pool, and diverge after a bottleneck point when they branch into dI-specific trajectories (Fig 1F: dI1: *Lhx9*^+^/*Barhl2*^+^; dI2: *Foxd3*^+^; dI3: *Isl1*^+^; dI4: *Pax2*^+^; dI5: *Lmx1b*^+^; dI6: *Dmrt3*^+^). The dI1 and dI5 lineages immediately emerge as distinct trajectories, while dI2, dI3, and dI4 initially share a common progenitor lineage before branching (Fig. 1D, F). We did not observe a distinct trajectory for dI6, rather it arises between the endpoints of the dI2 and dI4 lineages (Fig. 1D).

Unsupervised clustering of the dataset yields 31 clusters, which further subdivide the progenitor and dI lineages (Fig. 1E, G). The top 50 genes present in each cluster can be assessed in Table S1. The progenitor domain clusters are enriched for genes regulating the cell cycle, including S-phase (*Pcna*, *Mcm4, Gmnn*) (Komamura-Kohno et al., 2006; Kushwaha et al., 2016; Zerjatke et al., 2017), G2/M-phase (*Cdc20*, *Aurka*) (Cazales et al., 2005; Lara-Gonzalez et al., 2019) and G1-phase (*Ccnd1*) (Wang et al., 2018), as well as roof plate markers (*Msx1*) (Liu et al., 2004) (Fig. 1G). Clusters 25 and 11 span the transition from pan-dP to dI2/dI3/dI4 identities. Cluster 25 is enriched for broadly expressed dP markers including *Neurog1/2*, *Pax3* and *Olig3,* while cluster 11 shows expression of both dP markers and dI2/dI4 markers, such as *Lhx1/5* and *Pou4f1* (previously known as *Brn3a*) (Alaynick et al., 2011). Similarly, cluster 27 represents the first transition step of dPs towards the dI5 identity, expressing both dP5 markers, such as *Ccnd1* and *Ascl1*, and post-mitotic dI5 markers, like *Tlx3* and *Lbx1*. We see little to no expression of ventral (v) interneuron markers, such as *Pitx2*, *En1*, and *Gata2,* in this dataset confirming the dorsal specificity of the RA±BMP4 differentiation protocol (Fig. 1G).

### Pseudotime analysis identifies new transition state-specific markers for the dIs

To investigate the temporal changes in gene expression in the five differentiation trajectories (Fig. 1D), we performed pseudotemporal ordering using Monocle3 (Cao et al., 2019), with distance calculated based on the Sox2^+^ progenitor population as the starting root. The pseudotime values were then superimposed onto the UMAP atlas, to reveal the distance and trajectories over which the progenitors differentiate into post-mitotic neurons (Fig. 2A). After optimization (see methods), the monocle trajectories (Fig. 2A, Supplemental movie 1) reveal similar bifurcation points to the cluster-based trajectory assignments (Fig. 2B). First, the dI5 branch splits off from dI1-dI4 lineages, rapidly followed by the dI1 branch splitting from the dI2-dI4 lineages. The dI2/dI3/dI4 lineages continue on a common path until they bifurcate, first to yield dI4 vs. dI2/dI3, and then dI2 vs. dI3 (Fig. 2A, B). Interestingly, we also observed that the Monocle and lineage trajectories converge upon terminal differentiation (Fig 1C, Fig 2A, B, Supplemental movie 1). After branching to become distinct developmental trajectories, the dI2/dI3/dI4/dI5 lineages then merge as they differentiate as if they have again become more transcriptionally similar to each other. While the dI1 trajectory remains distinct, it also curves towards the same region of statistical similarity occupied by the other terminal branches.

**Figure 2:**
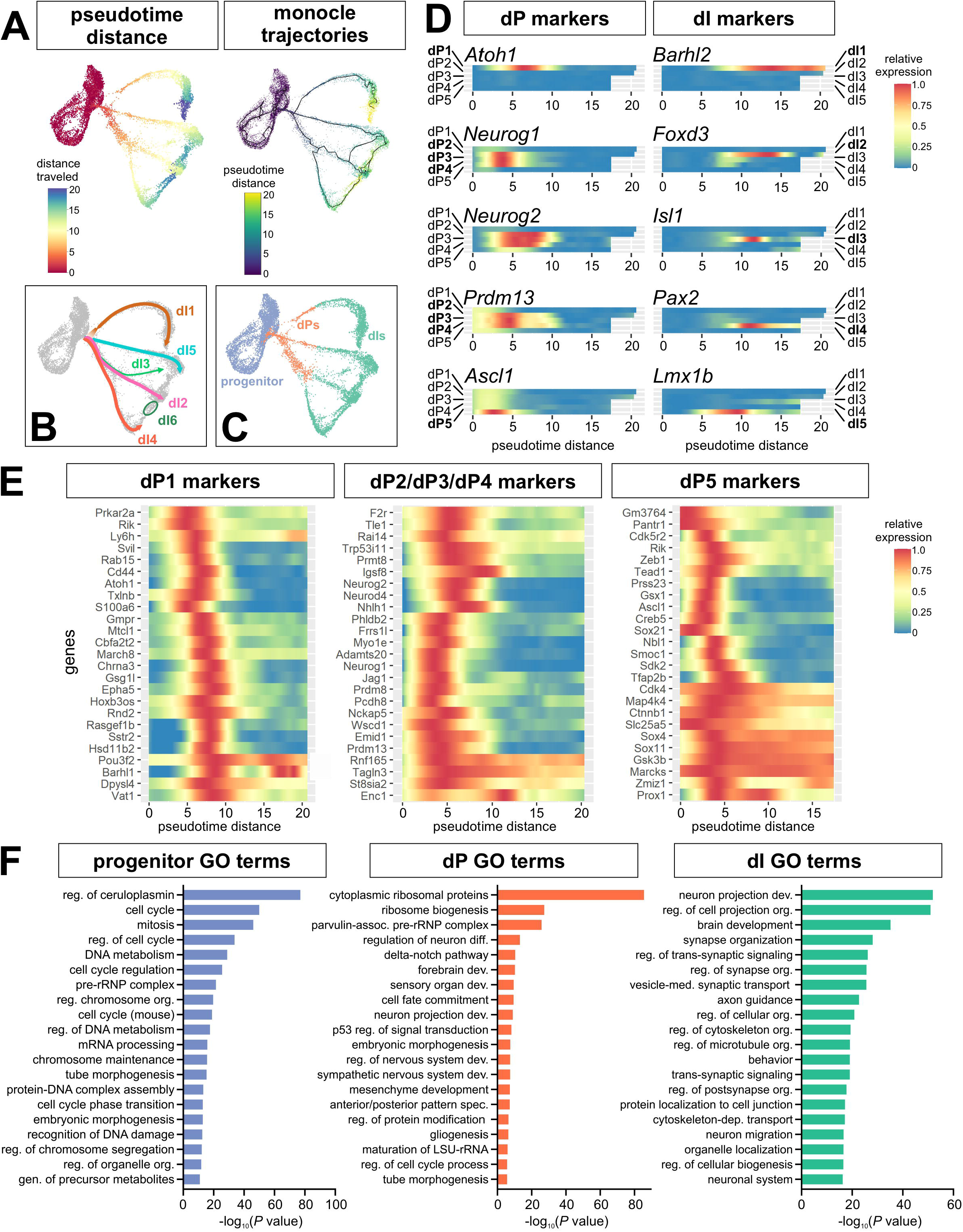
Using pseudotime to identify dI-specific trajectories and markers *in vitro*. (A) Pseudotime analysis identifies both the distance and trajectories over which progenitors transition to dPs and then differentiated dIs. (B) Putative dI trajectories based on the marker analysis in Fig. 1D. (C) Based on gene expression, we classified progenitors to be the root cells used in Monocle3, dPs as being from >0-7.5 pseudotime distance, and differentiated dIs from ≥7.5 – 20 pseudotime distance. (D) Heatmaps drawn from the dI1-dI5 pseudotime trajectories show the correct temporal distribution of cardinal dP and dI markers. (E) Heatmaps showing the expression of marker genes expressed in the three major dP clusters, i.e., dP1, dP2/dP3/dP4 and dP5, many of which are novel. (F) Gene ontology (GO) analysis shows enriched biological processes in the clusters assigned to the progenitor, dP, and differentiated dI identities, as shown in panel C.

To follow gene expression changes over pseudotime, we first examined the temporal distribution of known marker genes. In every case, we find that the canonical dP and dI markers have the correct lineage-specific and temporal expression patterns (Fig. 2D). Thus, dP markers start to be expressed prior to pseudotime values of 5, while the expression of dI markers tends to peak near or after pseudotime values of 10, suggesting that pseudotime distance accurately reflects developmental time *in vivo*. We next subdivided the data into progenitor, dP, and dI populations based on their pseudotime values and trajectory divergence points (Fig. 2C), to identify genes expressed in the different differentiation states, and performed gene ontology (GO) analyses for enriched terms (Fig 2F). While the progenitors and dIs showed the predicted enrichment of cell cycle related terms, and synapse formation related terms respectively, the dPs were enriched for terms related to neural differentiation, patterning and cell fate, further supporting the conclusion that dP clusters (between pseudotime value >0-7.5) represent transition states (Fig. 2F).

To identify candidate genes that establish these transition states for different dI trajectories, we performed differential gene expression on all clusters. We then compared the top marker genes from the first cluster in the dI1 (clusters 15) vs. dI2/dI3/dI4 (cluster 25) vs. dI5 (cluster 27) lineages, after removing any common markers appearing in multiple lists (Fig. 2E). We thereby identified genes showing enriched expression in different dI lineages. Gene expression was first validated using UMAP plots to determine the extent to which these genes were present in specific trajectories (Fig. S2). We found that the genes were generally expressed in the predicted trajectories, but were not always specific to that lineage. This analysis (Figs. 2D and S2) identified both canonical transcription factors known to be important for establishing dP fates, including *Atoh1* (dP1, (Helms and Johnson, 1998)), *Neurog1* (dP2), (Gowan et al., 2001) and *Neurog2* (dP2-dP5) (Sommer et al., 1996) validating our methodology, and many novel genes whose expression patterns were then assessed *in vivo*.

### Validation of putative transition state markers in vivo

Nine genes showing enriched expression in the dP transition clusters were selected for further analysis. The expression of these genes was examined in transverse sections of E10.5 and E11.5 spinal cord through *in situ* hybridization analyses (Fig. 3). Of these nine genes, one - *Gsg1l*, a known Atoh1 target gene (Lai et al., 2011), which encodes a regulatory subunit of AMPA receptor (Kamalova et al., 2021) – shows specific expression in the dI1 lineage (arrows, Fig. 3A) as predicted by the pseudotime analysis (heatmap, Fig. 3A; clusters 15/26, Fig. 1E). Two additional genes - *Chrna3,* which encodes a cholinergic receptor subunit (Flora et al., 2013), and *Cbfa2t2,* a transcriptional corepressor (Tu et al., 2016) *–* are initially broadly expressed by differentiating neurons in the E10.5 dorsal spinal cord (Fig. 3B, C). By E11.5, the expression of both genes becomes more prominent in the dP1/dI1 lineage and the intermediate zone (IZ), the region where differentiating dPs exit the ventricular zone (VZ) as they migrate laterally to become postmitotic neurons (Fig. 3B). Again, this distribution strikingly mirrors the predictions from the pseudotime analyses (heatmaps, Fig. 3B, C), especially for *Cbfa2t2,* which is expressed sequentially first in dP5 (cluster 17, Fig. 1E) and then dP1/dI1 (clusters 15/26, Fig. 1E), with lower expression in the rest of the dorsal IZ (arrows, Fig. 3C).

**Figure 3:**
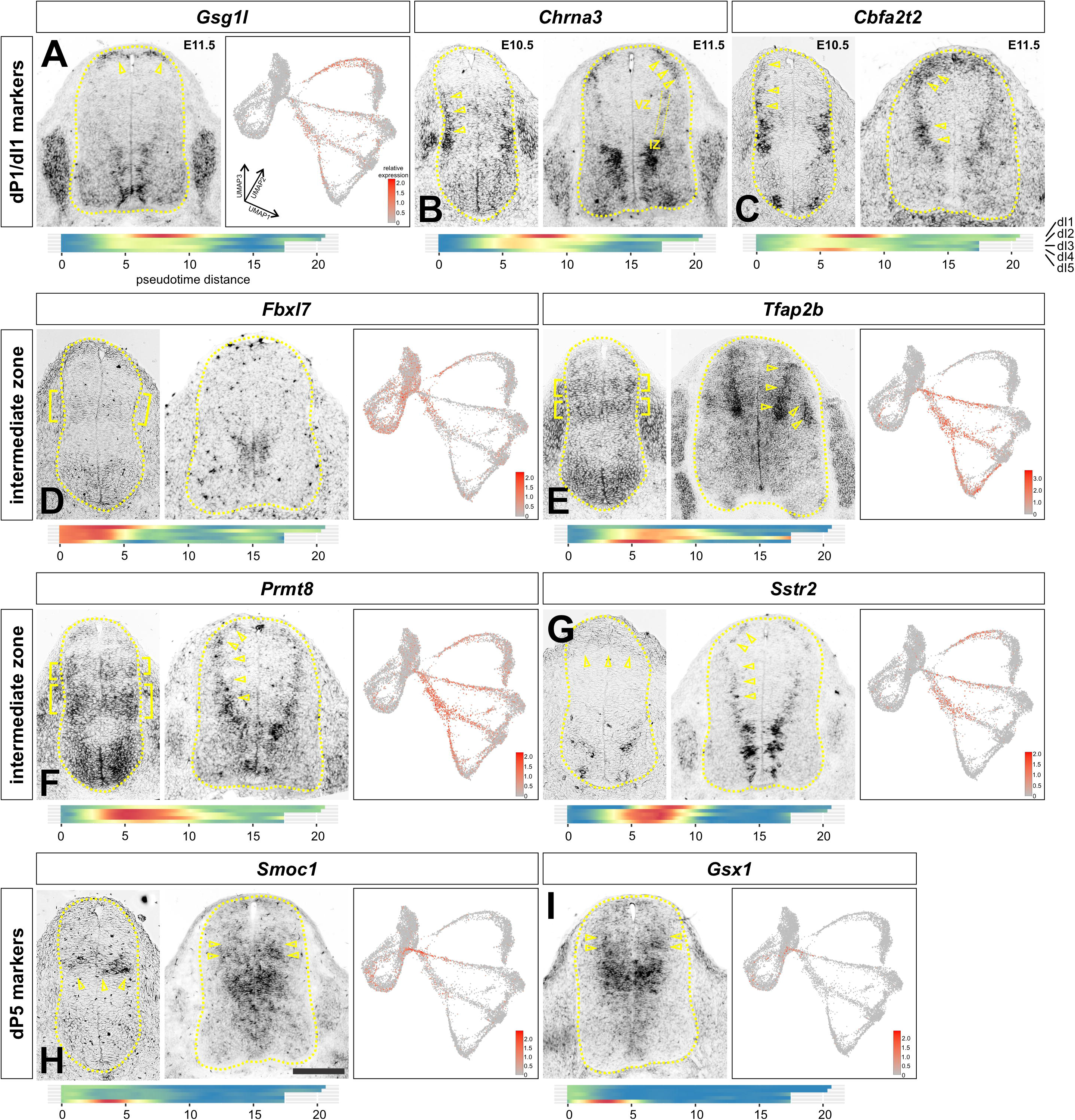
*In vivo* validation of dI lineage markers identified *in vitro*. Gene expression was assessed in E10.5 and E11.5 lumbar and thoracic spinal cord sections by *in situ* hybridization and compared to the predicted distribution and timing of expression from the UMAP reduction and pseudotime ordering (heatmap). (A) *Gsg1l* is specifically expressed in dP1s, both *in vivo* and *in vitro*. (B, C) *Chrna3* and *Cbfa2t2* are expressed in newly differentiating dIs, i.e., in the intermediate zone (IZ, dotted lines), with highest levels in the dI1s. The heatmap shows similar enriched expression in the transitory region of the dI trajectories. (D) *Fbxl7* expressed at low levels in intermediate dPs in the ventricular zone (VZ) at E10.5. Expression then diminishes by E11.5, as predicted *in vitro*. (E, F) *Tfap2b* and *Prmt8* are expressed in stripes in dPs at E10.5. Expression resolves to the newly differentiating dIs in the IZ at E11.5, as predicted by the heatmaps. (G) *Sstr2* is expressed in dorsal dPs at E10.5 and differentiating dIs in the IZ at E11.5, corresponding to the transitory clusters in the heatmap. (H-I) *Smoc1* and *Gsx1* are expressed broadly in dPs, with highest expression in the dP5 domain. Scale bar: E10.5 images: 50µm; E11.5 images 100 µm

Four of the selected genes *– Fbxl7, Tfap2b, Prmt8*, and *Sstr2* – are predicted to be present in subsets of multiple dPs based on the pseudotime analysis. Three of these genes - *Tfap2b,* an AP2 family transcription factor (Zainolabidin et al., 2017), *Prmt8,* an arginine methyltransferase (Dong et al., 2021), and *Sstr2*, somatostatin receptor 2 (Stumm et al., 2004) – show expression first in the VZ (E10.5) followed by robust increases in the IZ (E11.5) *in vivo* (arrows, Fig. 3E, F, G). The fourth gene - *Fbxl7,* part of the ubiquitin ligase complex, is expressed in the intermediate VZ, but not upregulated in the IZ (bracket, Fig. 3D) as predicted *in silico*. These genes are expressed in multiple dI lineages. For example, *Tfap2b* is upregulated in the IZ specifically in the dI2-dI5 lineages in both the *in vitr*o-derived atlas and in the E11.5 spinal cord (Fig. 3E). The remaining two genes - *Smoc1*, a secreted calcium-binding protein (Thomas et al., 2017), and *Gsx1*, a previously identified spinal cord transcription factor (Mizuguchi et al., 2006) - were also validated from the *in silico* data. Both genes were predicted to be expressed at the beginning of the dP5 trajectory (cluster 27, Fig. 1E), which was borne out in the *in viv*o analysis (arrows, Fig. 3H, I). In particular, *Smoc1* is specifically expressed in the dP5 domain in the VZ of both E10.5 and E11.5 spinal cord (arrows, Fig. 3H).

Finally, many of the nine *in silico*-identified genes - *Chrna3, Cbfa2t2, Tfap2b*, *Prmt8, Sstr2* - show striped expression patterns in the VZ and/or IZ, which is a hallmark of genes directing neurogenesis in a domain-restricted manner (Marklund et al., 2010; Skaggs et al., 2011). For example, both *Tfap2b* and *Prmt8* are expressed in two stripes in the VZ at E10.5 (brackets, Fig. 3E, F), while *Prtm8* and *Sstr2* are discontinuously expressed in the IZ (arrows, Fig. 3F, G). These genes are thus candidates for factors that direct multipotential progenitors into more restricted transition states before resolving into specific dI lineages.

### Comparison of in vivo and in vitro scRNA-Seq datasets

We next compared our *in vitro* atlas to an *in vivo* scRNA-Seq dataset of the embryonic spinal cord collected between E9.5-E13.5 (Delile et al., 2019). We identified the differentiation trajectories of the *in vivo* cells by isolating neural progenitors and neurons, splitting the dataset into different timepoints and then reintegrating the timepoints by the reciprocal PCA method (see methods). This approach yielded clear lineage trajectories for the different dorsal and ventral spinal neurons (Fig. 4B). In this dataset, the *Sox2*^+^ progenitor pool produces two major bottleneck points which respectively lead to the dorsal and ventral (v0-v3 and motor neurons (MNs)) trajectories (Fig. 4B, see https://samjbutler.shinyapps.io/BriscoeVisualization/ to assess the trajectories in three dimensions). In the dorsal trajectories, the dI1 lineage emerges as a distinct trajectory, while dI3, dI4 and dI5 initially share an extensive common progenitor lineage before branching into dI4 vs dI3/dI5, with dI3 subsequently branching off from dI5. The starting point of the dI2 trajectory is not clear, either branching from dI1, or originating from dI3/dI4/dI5 bottleneck (arrow, Fig. 4B). Interestingly, dI6 does not arise with the other dorsal cell types, but rather shares an initial pathway with Evx1^+^ v0 interneurons. Thus, the overall pattern of the *in vivo* trajectories is similar to the *in vitro* atlas (Fig. 4A), although there are some differences, most notable of which are the origin of the dI2s, and the dI6s.

**Figure 4:**
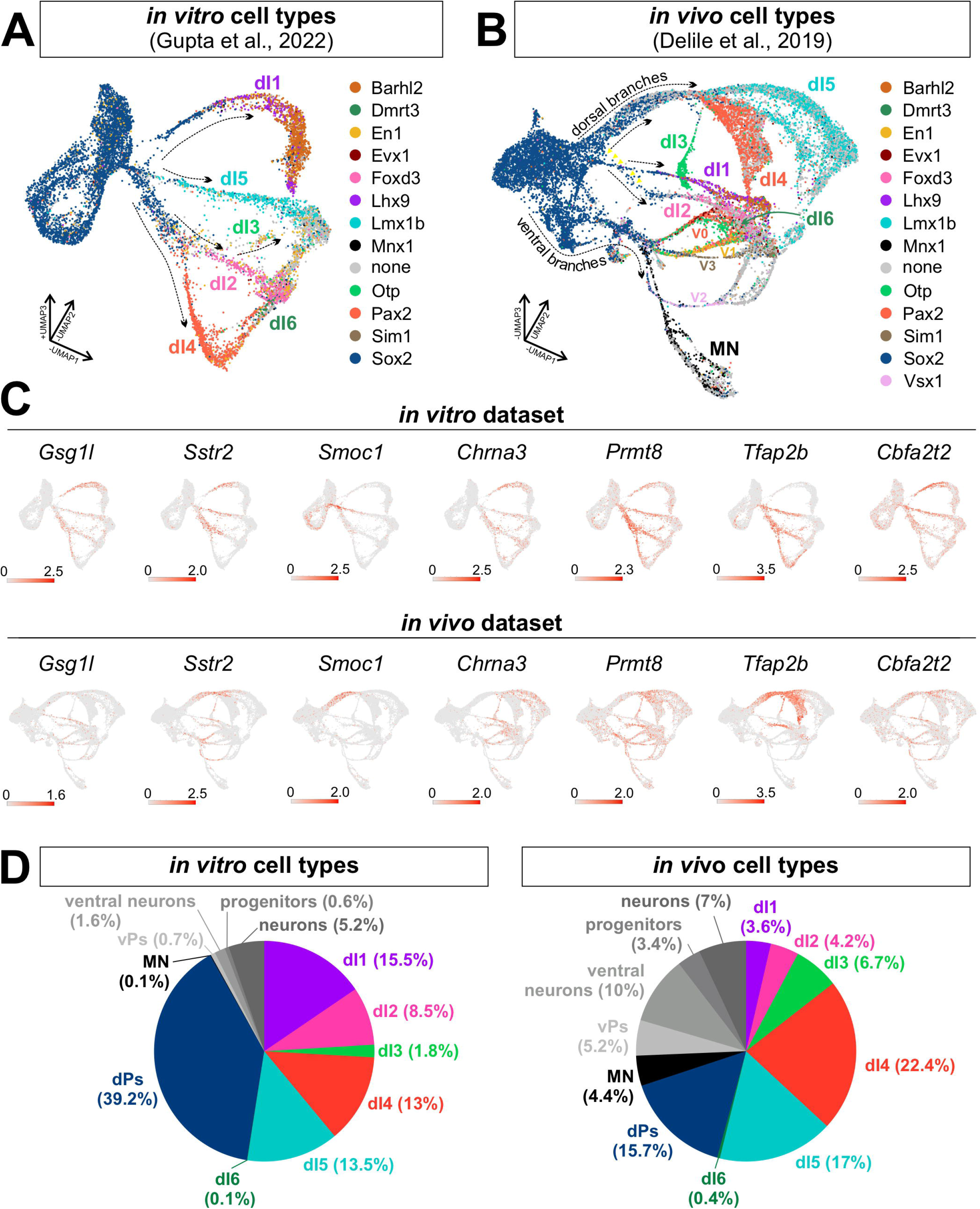
Comparison of in vitro and in vivo-derived dI trajectories. (A-B) UMAP feature plots of *in vitro* (A) and in vivo (B) spinal progenitors and neurons. The cell types were annotated according to (Delile et al., 2019). (C) UMAP feature plots showing that the putative “transition” genes” are generally expressed similarly in the *in vitro* and *in vivo* atlases. (D) Analysis of cellular proportions in the single cell atlases using the nearest neighbor method demonstrates that the *in vitro* dataset is enriched for dPs and dIs, rather than ventral progenutors (vPs) and neurons. A small percentage of progenitors and neurons could not be assigned to a specific cell type in each atlas.

This larger dataset also reveals the existence of multiple dI4 and dI5 subtypes. Both lineage trajectories split into two (dI4) or three (dI5) paths that align closely within the overall differentiation trajectory. However, after an initial period of expansion, the dI4/dI5 trajectories then converge back together towards other interneuron populations during the terminal differentiation process (Fig. 4B). Thus, as observed in the *in vitro* atlas (Fig, 4A), spinal neurons - with the exception of the dI3s and MNs - become markedly more transcriptionally similar as they mature. We then integrated the *in vitro* and *in vivo* datasets together, and found that ∼92% of the *in vitro* cells overlapped with dorsal identities *in vivo*, while ∼2% mapped to the ventral identities (Fig. 4F, Fig. S3). ∼98% of the *in vitro* progenitors corresponded to the (Delile et al., 2019) dorsal progenitor category, despite there being no discernable divisions in the Sox2^+^ progenitor zone in the UMAP (Fig. S3A).

We then further validated the *in vitro*-identified “transition” genes using the *in vivo* scRNA-Seq dataset (Fig. 4C). We generally saw striking conservation of the expression patterns, with the exception of *Fbxl7,* which was minimally expressed in the *in vivo* atlas (data not shown). However, *Fbxl7* also had the weakest signal in the *in situ* hybridization analysis (Fig, 3D), raising the possibility that the read depth was not sufficient for detection. The other eight genes were expressed in the same dorsal lineage trajectories both in *in vivo* and *in vitro* (Fig, 4D). *Prmt8*, *Cbfa2t2*, *Chrna3* and *Sstr2* were additionally expressed in the ventral identities predicted by the *in situ* hybridization analyses. However, *Gsg1l* and *Smoc1* showed minimal expression in ventral cell types *in silico* (Fig. 4D), despite visible expression *in vivo* (Fig. 3A, 3H), further highlighting possible sensitivity differences between the validation techniques.

### Pseudotime analysis reveals subtype diversification of dIs

While many clusters could be identified using canonical markers, the identity of cluster 16 could not be resolved (Fig. 1E, 5A). By eye, cluster 16 is consistent with being at the endpoint of the *Foxd3^+^*dI2 lineage (Fig. 1D), however, cluster 16 is derived from the RA protocol which generates mostly dI4/dI5/dI6s (Fig. 1C). Supporting this possibility, the Monocle trajectories suggested that either the dI5 or dI4 lineages could contribute to cluster 16 (Fig. 5B). To resolve the identity and origin of cluster 16, marker gene analysis was used to identify *Sncg* - synuclein gamma - as the most significantly upregulated gene in cluster 16 (Fig. 5C, 5D). GO analysis of the *Sncg*^+^ expressing cells vs. those exclusively in cluster 16 showed similar enrichment of terms related to axon projection and synapses, indicating that these cells may collectively participate in establishing long range connections (Fig. 5F, 5G).

**Figure 5:**
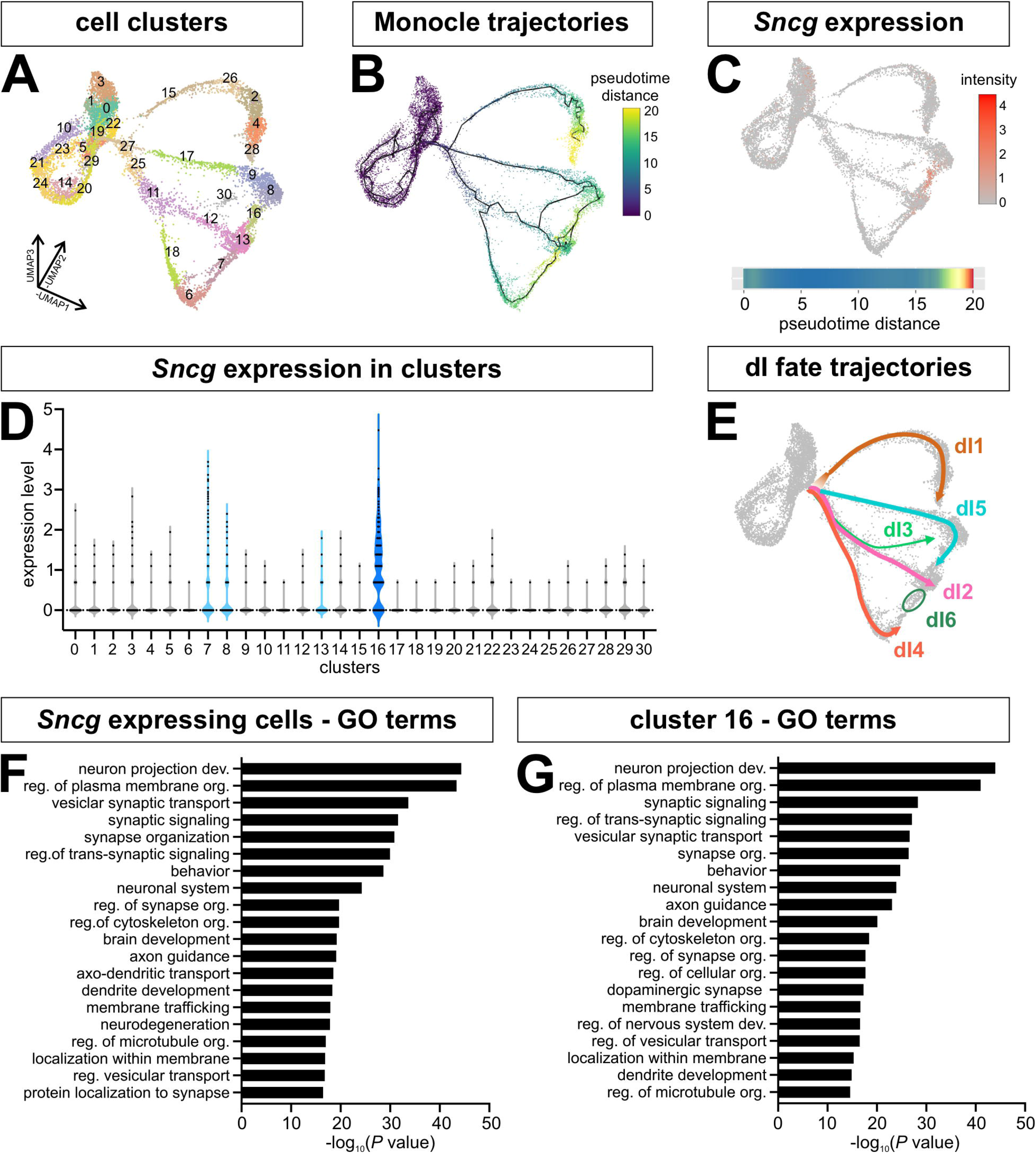
Characterization of Synuclein-γ (Sncg)^+^ cluster 16. (A) Unsupervised clustering resolves the dI trajectories into 31 clusters. (B) The Monocle3 derived pseudotime trajectories suggest that dI5, dI4 and dI2 could converge on cluster 16. (C-D) Synuclein-γ (Sncg) is most robustly expressed in cluster 16, with some expression in cluster 7, an adjacent cluster. Expression in the other two adjacent clusters, clusters 8 and 13, is not above background. (E) The dI fate trajectories were redrawn from Fig.1D, 2B to depict cluster 16 as the end point of the dI5 lineage. (F, G) Similar GO terms, related to axon guidance and synapse formation, were enriched in the Sncg expressing cells (F) and cluster 16 (G).

To further investigate these populations, we performed a marker gene analysis on *Sncg* positive cells in the terminal region (clusters 16 and 7). This analysis identified *synaptotagmin* (*Syt*) *4* and *Syt13* (Fig. S4A) which are expressed in the Phox2a^+^ dI5 subtype that relays pain and itch to the thalamus (Roome et al., 2020), suggesting that cluster 16 is at the endpoint of the dI5 lineage (Fig. 5E). We further explored this hypothesis by analyzing *Sncg* expression in the E11.5, E12.5 and E13.5 mouse spinal cord *in vivo*, in combination with immunohistochemistry for Pax2, which labels dI4 and dI6 (Gross et al., 2002), and Lmx1b, which decorates dI5s (Ding et al., 2004) (Fig. 6). At both stage E11.5 and E12.5, *Sncg* expression colocalizes with Lmx1b^+^, but not Pax2^+^, cells (inset Fig. 6A, 6B), strongly suggesting that cluster 16 represents a dI5 subtype (Fig. 5E). The anatomical position of this *Sncg^+^*Lmx1b^+^ cluster is also consistent with the Phox2a^+^ dI5 population identified by lineage tracing (Roome et al., 2020). However, Phox2a is not robustly expressed in cluster 16 (Fig. S4B), and the co-localization of *Sncg* transcripts within Lmxb1^+^ cells does not persist into stage E13.5 (inset, Fig 6B, 4C). We thus additionally assessed the distribution of Sncg by immunohistochemistry at E12.5 (Fig. 6D) and E13.5 (Fig. 6E). This analysis demonstrated that the dorsal-most population of Sncg^+^ neurons continues to be Lmx1b^+^ (inset, arrows, Fig. 6D, 6E). Moreover, these Lmx1b^+^ Sncg^+^ neurons appear to be migrating dorsally by E13.5, similar to previous observations showing that Phox2a^+^ dI5s migrate tangentially to the upper laminae of the dorsal horn starting at E13.5 (Roome et al., 2020). By postnatal day 4, *Sncg*^+^ cells are present on the surface of the dorsal horn in a position consistent with a subset of dI5s (Fig. S4D). It remains unresolved whether the dorsal-most cells that continue to express *Sncg^+^*at E13.5 (inset, Fig. 6C, inset) represents a previously undescribed class of dI5s, or more likely a subset of MNs.

**Figure 6:**
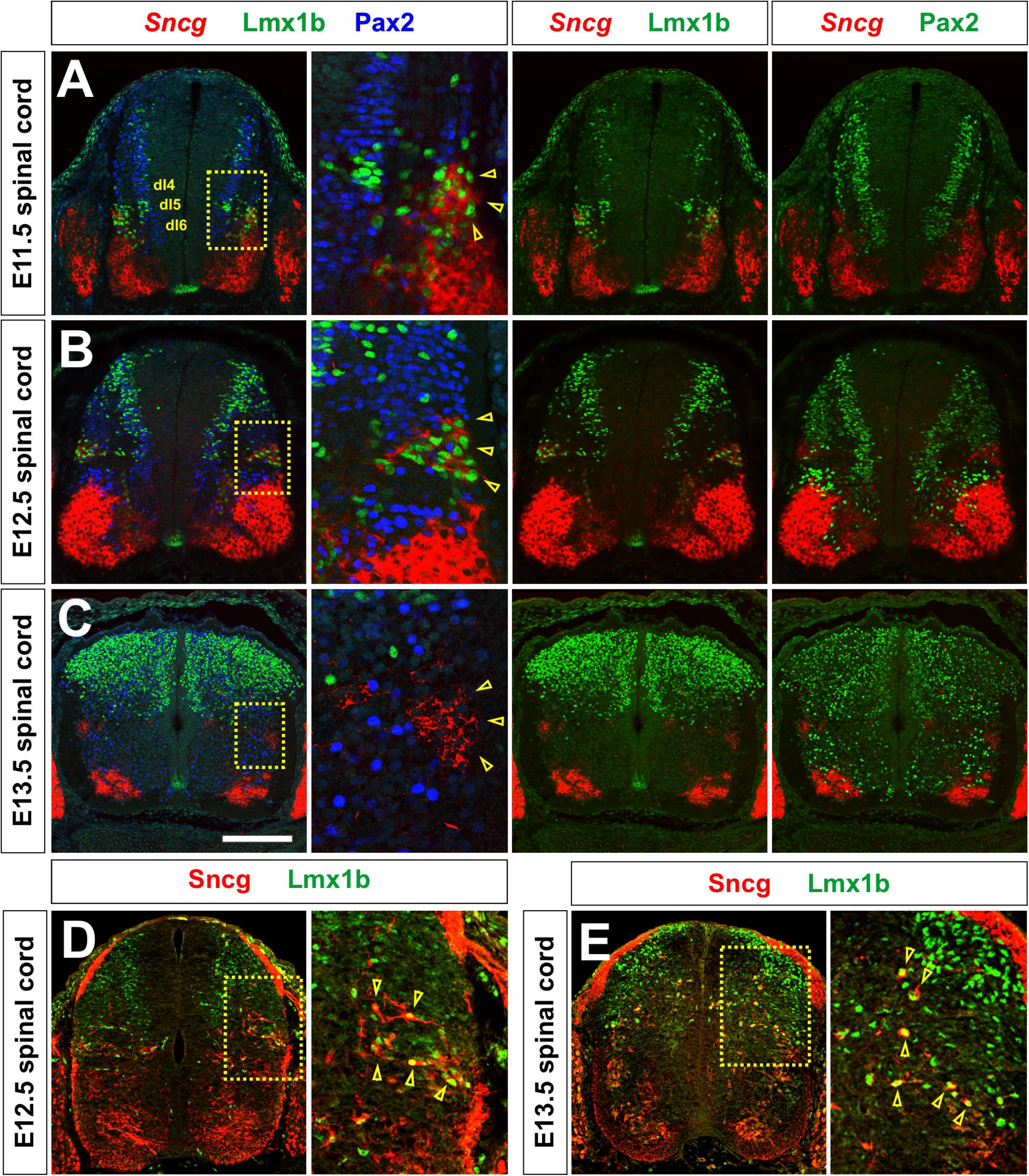
Identification of *Sncg*-expressing cells as a dI5 subtype. Thoracic (A-C) or lumber (D, E) transverse sections from E11.5 (A), E12.5 (B, D) and E13.5 (C, E) mouse spinal cords were subjected to both *in situ* hybridization to detect *Sncg (*red, A-C) and/or immunohistochemistry to detect Sncg (red, D, E), Pax2 (dI4 blue, green) and Lmx1b (dI5, green). (A, B) At both E11.5 and E12.5, *Sncg* expression is observed in a cluster of Lmx1b^+^ cells (inset), and not in Pax2^+^ neurons, in the dorsal spinal cord. The lateral position of *Sncg-*expressing cells is coincident with the Phox2a^+^ dI5 subtype (Roome et al., 2020). (C) By E13.5, the dorsal-most population of *Sncg^+^*cells no longer express Lmx1b, or Pax2 (inset). (D-E) However, a subset of Lmx1b^+^ cells continue to be Sncg^+^ at both E12.5 (D) and E13.5 (E), and appear to be migrating towards a more superficial layer of the spinal cord. Scale bar: (A, B) 150µm, (C) 200µm

### Online tool to investigate gene expression dynamics during dI differentiation in vitro

We have created two online data tools to visualize the distribution of genes present in the UMAP reduction format both *in vitro* (https://samjbutler.shinyapps.io/Data_Viewer/) and *in vivo* (https://samjbutler.shinyapps.io/BriscoeVisualization/), together with the pseudotime trajectory heatmaps (Supplemental movie 2). A variety of parameters can be adjusted using the menus including the span size for fitting a smooth curve, i.e., LOESS regression, and size of the plots. This tool provides a starting point for researchers to discover new dI-associated genes in our dataset, by visualizing their expression patterns and temporal changes, and thereby inferring function. This resource, and its associated data sets, may further serve as a roadmap for the derivation of dIs from both mouse and human ESCs and iPSCs and enable functional comparisons across species.

## Discussion

We have established a tool to track dI fate specification both spatially and temporally *in silico* by adding pseudotemporal ordering to our scRNA-Seq atlas of the developing dorsal spinal cord. We have used this tool to identify the lineage relationships between different trajectories and predict the identity of novel regulators of dI fates. Our *in vivo* data validate this approach and further illustrate how the RA±BMP4 directed differentiation protocols can be used to investigate dI cell fate decisions. We anticipate that this tool, together with an *in vivo* atlas constructed from the (Delile et al., 2019) dataset, will serve as valuable resources to explore the genes and molecular pathways that contribute to the development of the somatosensory system in the spinal cord.

### A spatial and temporal map of dI differentiation

Our analyses (Fig. 5E) distinguish five distinct trajectories corresponding to the dI1-dI5 populations. In contrast, dI6s emerge at the end of the dI4 trajectory (Fig. 1D). This observation may stem from the inefficient generation of dI6s *in vitro*, resulting in too few cells to delineate the dI6 lineage trajectory. Alternatively, it may rather accurately reflect endogenous development in the spinal cord, i.e., that dI6s arise from a common progenitor pool with dI4s and become specialized post-mitotically. Supporting the latter possibility, dI6s and dI4s express many of the same transcription factors, including *Pax2*, *Lbx1* and *Lhx1/5* (Lai et al., 2016). An additional possibility is that the dI6 population arises from both dorsal and ventral progenitors. Previous studies have suggested that dI6s *in vivo* arise in part from more ventrally located p0 progenitors (Griener et al., 2017). This hypothesis is supported by the *in vivo* atlas (Fig. 4B) where dI6 shares an initial pathway with v0, before diverging as they mature. The ventral lineage of dI6 might explain why our dorsal directed differentiation protocols produce so few dI6s. However, it is also notable that the dI6 population is smaller than the other classes of dIs both *in vitro* (∼0.1%,) and *in vivo* (∼0.4) (Fig. 4D), suggesting that the number of dIs produced may be an intrinsic feature of these dP populations.

The *in vitro* atlas also suggests a lineage relationship between the dI2 and dI4 populations, which have some common elements to their transcriptional code, i.e., *Neurog2^+^*dPs which resolve into *Lhx1/5^+^* dIs (Lai et al., 2016). *In vitro*, both dI2s and dI4s arise from the same dP lineage, which branches when dPs express either *Foxd3* (dI2) or *Pax2* (dI4) (Fig. 1D, Fig. 2A-C). This shared relationship may explain why the RA protocol makes small numbers of dI2s, while the RA+BMP4 protocol has a small dI4 population (Gupta et al., 2022). However, we were unable to unambiguously validate this dI lineage relationship in the *in vivo* dataset (Fig. 4B). The *in vivo* data support two possibilities, that the dI2 lineage emerges either from the dI1 trajectory, or alternatively, from the shared dI3/dI4/dI5 state, with the dI2 lineage then crossing the dI1 trajectory (arrows, Fig 4B). Interestingly, while the *in vivo* and *in vitro*-derived dI types completely overlapped when the two data sets were merged (Fig. S3), the lineage relationships shifted, such dI2 lineage now again emerged from the dI4 trajectory (Fig S3D, S3E). However, an important caveat with these *in silico* studies is that they must be considered predictive. The correct lineage relationships can be further resolved by studies *in vivo*, such as lineage tracing with genetic barcodes or RNA velocity analyses. A discontinuous lineage relationship between dIs has been observed experimentally in *in vivo*: BMP7 depletion was shown to reduce the number of dI1, dI3 and dI5s in mouse (Le Dreau et al., 2012) while BMP4 overexpression in chicken dramatically increases the number of dI2s, potentially at the expense of dI4/dI6s (Andrews et al., 2017).

Our analysis of undifferentiated progenitors also reveals novel genes that may regulate dP fate specification. For example, *insulin growth factor binding protein (Igfbp) 5,* but not *Igfbp3*, is expressed in cycling spinal progenitors (Fig. 1G), suggesting they respond selectively to insulin-like growth factor signaling. Insulin signaling, acting though *Igfbp5,* regulates proliferation in other progenitor populations, such the olfactory bulb subventricular zone (Vicario-Abejón et al., 2003) and smooth muscle progenitors (Ahmad et al., 2020; Ren et al., 2008). Thus, modulation of insulin signaling could expand specific dP populations.

### dI transcriptional identities converge during terminal differentiation

Both the UMAP and pseudotime analyses suggest that dI lineage trajectories are at their most divergent during the period when dIs are assuming their unique fates. However, at more mature pseudotime values, the dI2-dI6s converge back together into a series of clusters (clusters 6, 7, 8, 13 and 16, Fig. 5A) with similar transcriptomes (Fig.5B, E). While there are some genetic differences between these clusters, their close proximity in UMAP space suggests that these dIs have become more transcriptionally homogenous. This convergence is also seen in the *in vivo* dataset for every neural cell type, except the MNs and dI3s, suggesting that transcriptomic convergence may be a general property of most spinal cord interneurons. One possibility to explain this observation is that significant differences in gene expression are required to direct dI cell fate specification, migration, and axon guidance. However, once dIs have completed the differentiation process, they then require lower transcriptional heterogeneity because they function similarly to each other. This phenomenon has also been observed during the development of dorsal root ganglia (Faure et al., 2020), oligodendrocytes (Marques et al., 2018), *Drosophila* brain (Michki et al., 2021) and the *C. elegans* nervous system (Hobert and Kratsios, 2019). Thus, the loss of transcriptional heterogeneity upon maturation may be a common theme for neural differentiation and may be used as a mechanism to induce diverse subtypes with shared functional identities (Osseward et al., 2021).

### Identification of novel regulators of dI differentiation

Our *in silico* analysis of *in vitro*-derived dI trajectories has identified novel putative regulatory genes including ion channels (*Gsg1l, Cacna2d1*), receptors (*Chrna3, Sstr2, EphA5*), actin binding proteins (*Svil, Mtcl1, Tagln3*), enzymes (*Prmt8, Fbxl7*) and transcriptional factors (*Tfap2b, Zeb1, Ted1*). Like previously identified genes that direct transitory states in dPs, i.e., *Neurog1*, *Neurog2* and *Ascl1,* several of the *in silico* identified genes such as *Chrna3*, *Prmt8*, *Cbfa2t2*, and *Tfap2b,* are expressed in stripes of progenitors during E10.5 spinal cord patterning (Fig. 3B. C, E, F). By E11.5, these genes, together with *Sstr2* (Fig. 3G) shift their expression to the IZ, i.e., in dPs that are exiting the cell cycle and differentiating into post-mitotic neurons. Taken together, these analyses suggest these factors regulate dI patterning and neurogenesis. Previous studies support this hypothesis. Both *Chrna3* and *Sstr2* are part of a *NeuroD1*-interacting network, during *NeuroD1* mediated reprograming of astrocytes to neurons (Ma et al., 2022). *NeuroD1* is also expressed in the IZ, with *Chrna3* and *Sstr2* (Brohl et al., 2008). Similarly, there is evidence of regulatory interactions between *Tfap2b* and *Ptf1a* in other neuronal contexts. For example, *Tfap2b* expression is regulated by *Ptf1a* during cerebellar and retinal neurogenesis (Jin et al., 2015; Zainolabidin et al., 2017). In the spinal cord, *Ptf1a* is expressed specifically in the dP4 domain, suggesting it may activate *Tfap2b* to regulate dI4/dI6 differentiation. *Tfap2b* may more generally regulate the differentiation of GABAergic neurons, since it continues to be expressed in the inhibitory neurons in the E18.5 spinal cord (Wildner et al., 2013).

### Subtype diversification of dIs in vitro

Our analysis of the terminally differentiated clusters has identified *Sncg*, as a new marker for a distinct dI5 subtype. Previously, *Sncg* expression was reported to be present in spinal MNs and dorsal root ganglia (Ninkina et al., 2003). The lateral dI5 cluster of *Sncg^+^* cells in the developing spinal cord initially coincides with the Phox2a^+^ dI5 subtype, which contributes to the anterolateral system (ALS) relaying pain and itch to the thalamus (Price and Dubner, 1977; Roome et al., 2020). By E13.5, lineage tracing has shown that Phox2a^+^ dI5s migrate away from this lateral position along a tangential route, and ultimately populate the superficial layer of the dorsal horn (Roome et al., 2020). A subpopulation of Sncg^+^ cells appears to follow this route. We observe the dorsal-most Sncg^+^ Lmxb1^+^ cells migrating to a more dorsal position by E13.5 (arrows, Fig 6E). While the migrating cells appear to have downregulated the transcription of *Sncg*, a cluster of cells expressing *Sncg* persists in the lateral position at E13.5 (inset, Fig. 6C). These cells may be a novel dI5 subtype, which will migrate dorsally at a later time point. However, we think it more likely that they are a subtype of MNs, given that *Sncg* expression resolves into more limited domains in the motor column by E13.5 (Fig. 6C). Interestingly, very few cells expressed *Phox2a* in our dataset (Fig. S4B). This discrepancy may reflect either the transient expression of *Phox2a* in dI5s *in vivo* (Roome et al., 2020), or the limited total number of Phox2a^+^ cells, which then was challenging to capture *in vitro*.

Our molecular analysis of the *Sncg^+^* population in our scRNA-Seq atlas also suggests a mechanism for dI diversification. Among the most highly expressed genes in the *Sncg*^+^ cluster are *D930028M14Rik, Onecut1*, *Onecut2, and Onecut3*. *Onecut* genes have been shown to represses the expression of *Pou2f2*, and thereby regulate the distribution of the dI2-dI6 populations (Masgutova et al., 2019). *D930028M14Rik* is also antisense to *Pou2f2* (O’Leary et al., 2016), suggesting it could contribute to *Pou2f2* repression. The downregulation of *Pou2f2* via *Onecut1/2* and *D930028M14Rik,* could represent a genetic switch allowing the *Sncg* population to diverge from the Phox2a^+^ dI5 population. Taken together, this analysis reveals that the *in vitro*-derived dIs diversify into subtypes also found *in vivo*, underscoring how closely the RA±BMP4 directed differentiation protocols recapitulate endogenous programs of dI cell fate specification.

## Acknowledgements

We would like to thank James Briscoe, Riki Kawaguchi, and members of the Butler and Novitch labs for discussions.

## Data Accessibility

Sequencing data is available at Gene Expression Omnibus accession number GSE185891 (Samples GSM5625332 and GSM5625333). Processed data and code available on request.

