## Supplemental Figures for "Investigating the basis of lineage decisions and developmental trajectories in the dorsal spinal cord through pseudotime analyses"

**Figure S1:** Inclusion and exclusion criteria for cells included in the trajectory analyses.

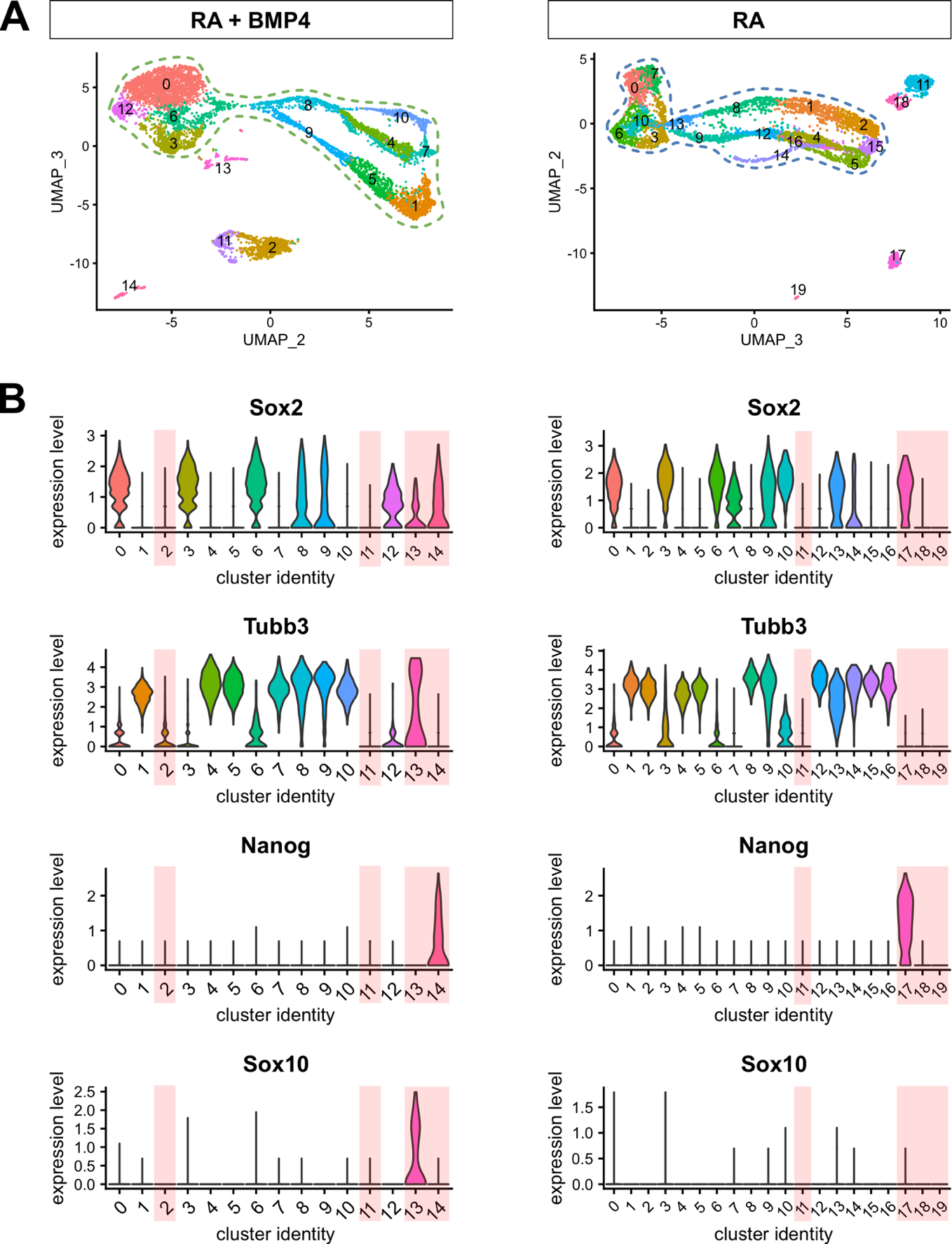

(A) UMAPs displaying the major cell clusters present in cultures resulting from the RA+BMP4 and RA-only protocols. Dashed lines illustrate cells that displayed neural progenitor or neuronal characteristics and were thereby included in the subsequent developmental trajectory analyses.

(B) Violin plots of gene expression in each cluster. Clusters (pink shading) that showed either: a) no *Sox2* or *Tubb3* expression or b) expression of *Nanog,* a pluripotent stem cell marker, or *Sox10*, a neural crest progenitor marker, were excluded from the datasets.

**Figure S2:** UMAPs of novel genes expressed in the three major dP clusters: dP1, dP2/dP3/dP4 and dP5.

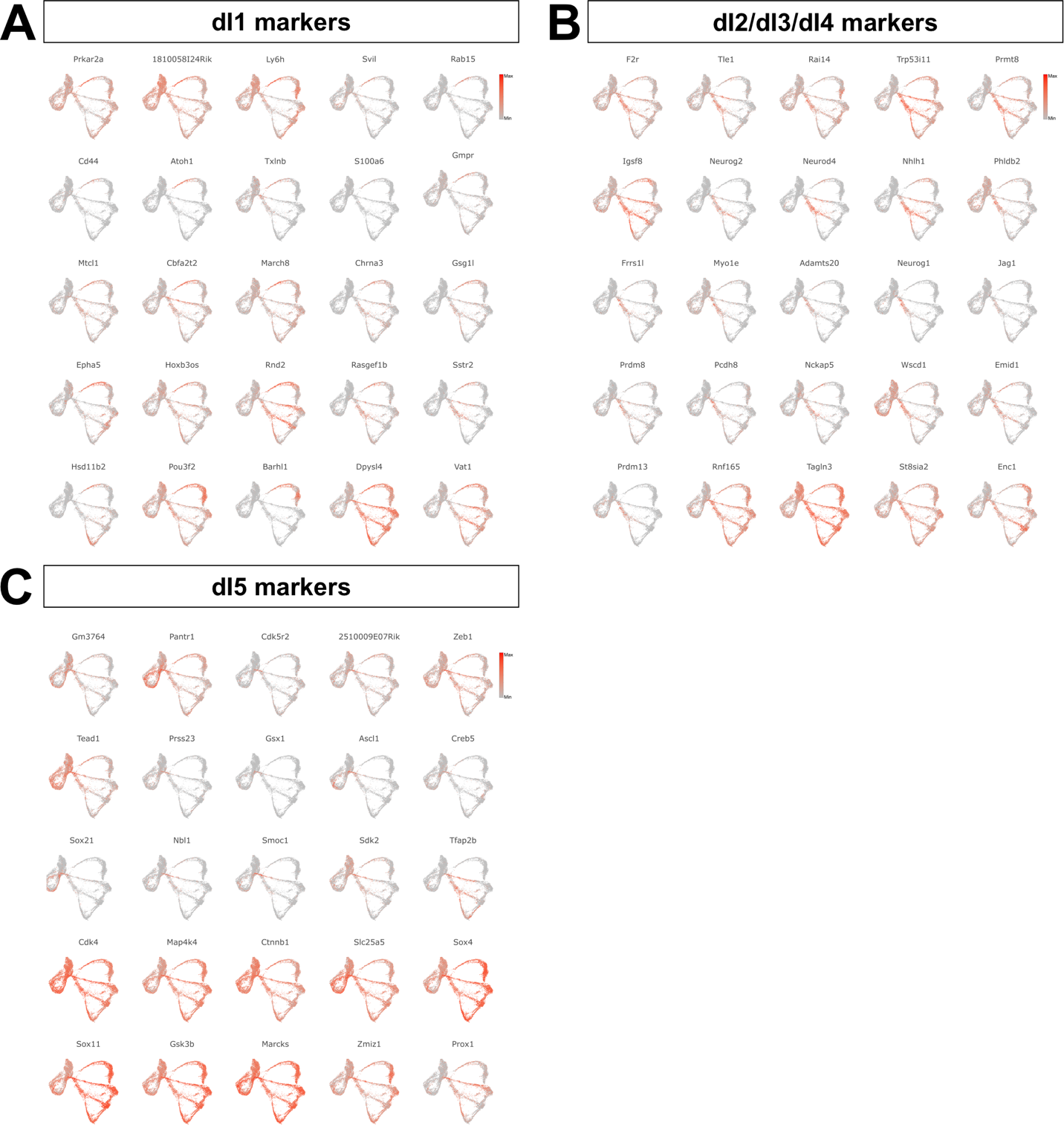

(A-C) UMAPs displaying the relative gene expression (minimal to maximal) of different genes that were enriched during the differentiation of dPs into dIs, as outlined in Fig. 2E.

**Figure S3:** Integration of the *in vivo* and *in vitro* derived scRNA-Seq datasets.

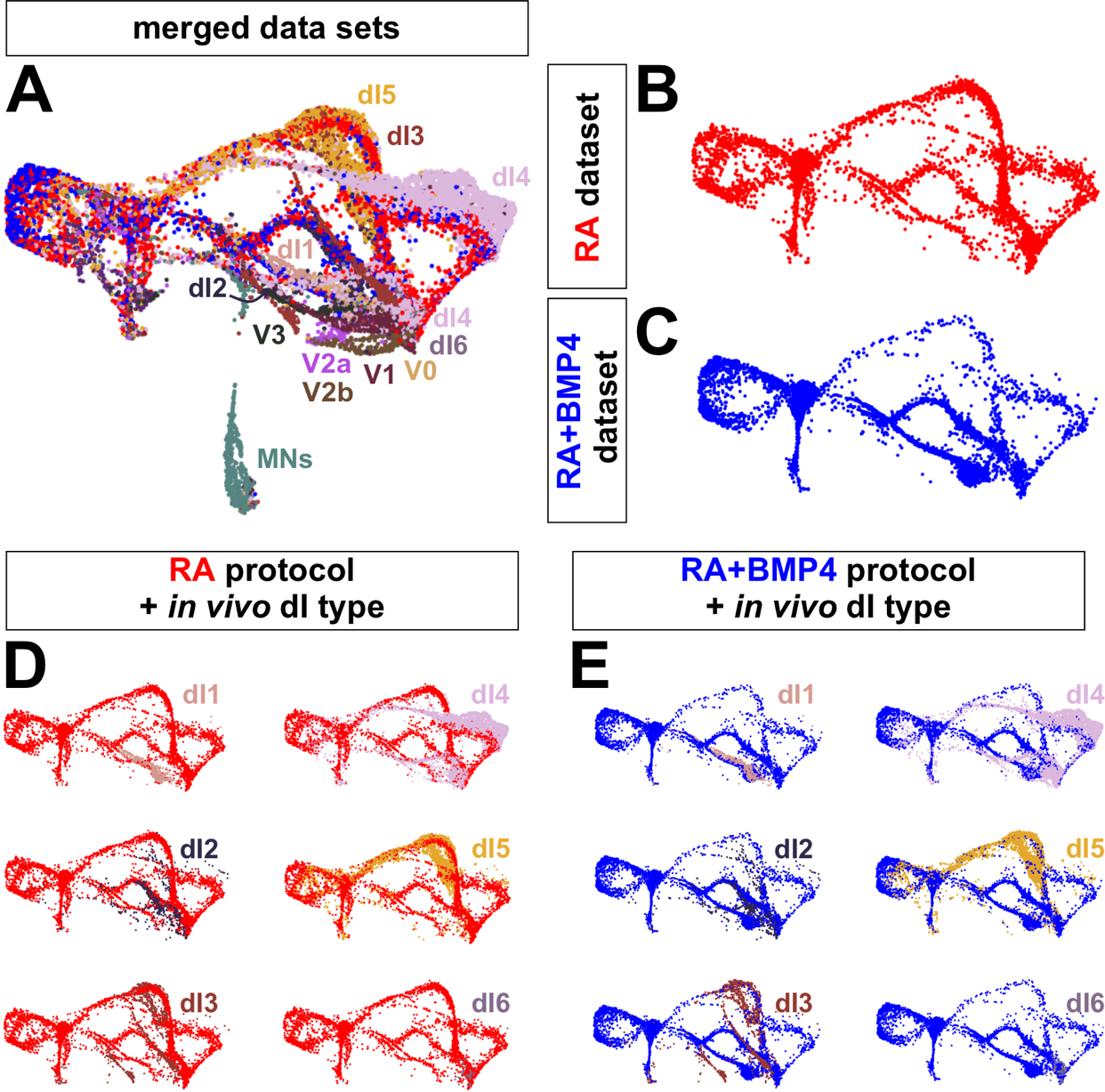

(A-C) The scRNA-Seq datasets from the developing spinal cord (Delile et al., 2019), and *in* *vitro-*derived dIs (Gupta et al., 2022) substantially overlap (A). While the UMAPs of RA dataset (B) and RA+BMP4 dataset (C) appear similar, there are key differences in the proportion of different dI trajectories, again demonstrating that these two directed differentiation protocols derive different subsets of dIs.

(D-E) Mapping individual *in vivo* dI types onto the RA (D) and RA+BMP4 (E) datasets show similar developmental trajectories, further suggesting that *in vitro* dI trajectories recapitulate embryonic spinal cord transcriptomic program.

**Figure S4:** Assessing the identity of cluster 16

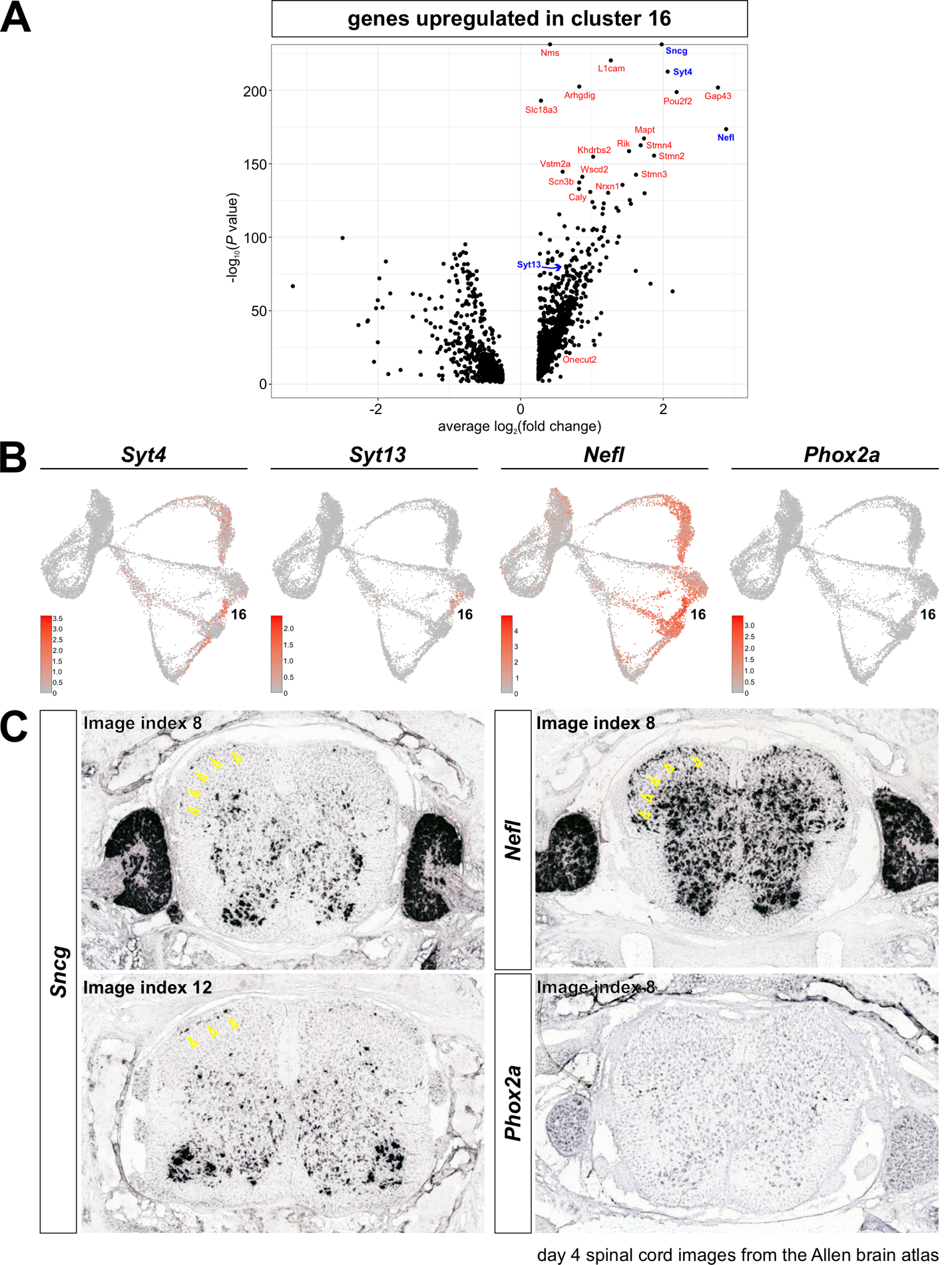

(A) Volcano plot showing the genes upregulated in cluster 16. These genes include *Syt4* and *Syt13* which have previously been identified as upregulated in the dI5 subtype (Roome et al., 2020)

(B) UMAP projections of genes associated with the thalamus innervating (ALS) dI5 subtype. These genes are also upregulated in *Sncg^+^* cluster 16, with the exception of *Phox2a.*

(C) *In situ* hybridization images of postnatal day 4 spinal cord, taken from the Allen brain atlas. Two cluster 16 markers - *Sncg* and *Nefl* – are present in cells on the surface of the dorsal horn (arrows). The location of these cells is consistent with that of the ALS dI5s. Note that *Phox2a* is only transiently present in ALS dI5 subtypes during development, and is thus not expressed in postnatal spinal cord.

**Supplemental Table 1:** The top 50 genes present in each cluster.

Attached as an Excel file.

**Supplemental Movie 1**

Supplementary movie showing Monocle3 trajectories. At later pseudotime values, the trajectories merge back together, with endpoints in clusters 16 and 7, which both express high levels of *Sncg.*

**Supplemental Movie 2**

Supplementary movie illustrating how the interactive tool can be used to visualize clusters, pseudotime distance, and expression patterns in the dataset. Multiple genes can be followed at once, to compare the relative intensity of gene expression.
